# Neuraminidase inhibitors — is it time to call it a day?

**DOI:** 10.1101/245175

**Authors:** César Parra-Rojas, Van Kinh Nguyen, Gustavo Hernández-Mejía, Esteban A. Hernández-Vargas

## Abstract

Stockpiling neuraminidase inhibitors (NAIs) such as oseltamivir and zanamivir is part of a global effort to be prepared for an influenza pandemic. However, the contribution of NAIs for treatment and prevention of influenza and its complications is largely debatable. Here, we developed a transparent mathematical modelling setting to analyse the impact of NAIs on influenza disease at within-host and population level. Analytical and simulation results indicate that even assuming unrealistically high efficacies for NAIs, drug intake starting on the onset of symptoms has a negligible effect on an individual's viral load and symptoms score. Increasing NAIs doses does not provide a better outcome as is generally believed. Considering Tamiflu's pandemic regimen for prophylaxis, different multiscale simulation scenarios reveal modest reductions in epidemic size despite high investments in stockpiling. Our results question the use of NAIs in general to treat influenza as well as the respective stockpiling by regulatory authorities.

## Introduction

Influenza A virus (IAV) infection affects about 20% of the worldwide population every year (***Moscona, 2005***). The 2009 influenza pandemic once again showed that the next pandemic will most likely result in major adverse health and economic outcomes (***Stöhr, 2005; Gates, 2015***). Although vaccination remains the primary means of control against outbreaks, vaccine developments for influenza are typically outpaced by the fast antigenic drift of the virus (***Ghedin et al., 2009***). In this scenario, the World Health Organization (WHO) has encouraged stockpiling antiviral drugs in anticipation of a pandemic (***Kelly and Cowling, 2017; Patel and Gorman, 2009***). The benefits of this approach, however, have been heavily debated, both in terms of treating and preventing epidemic spread (***Kelly and Cowling, 2017; Jefferson et al., 2014***).

Currently, the recommended antiviral drugs against influenza are neuraminidase inhibitors (NAIs) (***WHO, 2016***). NAIs block the release of influenza virus from infected host cells and could reduce the spread of infection in the respiratory tract (***Gubareva et al., 2000; Jefferson et al., 2014***). The drugs are generally safe to use for high-risk populations with risks of only mild adverse effects ***Jefferson et al., 2014; McClellan and Perry, 2001***). As the influenza infection course is fast, the effectiveness of NAIs depends strongly on the timing of the antiviral intakes, and their performance can be further compromised by the emergence of drug-resistant viruses (***Reece, 2007; Sheu et al., 2008***).

Clinicial trials found that oseltamivir—the most common NAI, sold under the brand name Tamiflu^®^—reduces viral shedding, lessens the disease severity, and shortens its duration by 1.5 days (***McClellan and Perry, 2001***). In adult subjects, oseltamivir reduced the time to first alleviation of symptoms of influenza-like illness by 16.8h ***Jefferson et al., 2014***). Treatment with NAIs could reduce mortality up to one-fifth, compared to the case without treatment (***Muthuri et al., 2014***). Early administration, within 2 days of symptom onset, reduced the mortality risk compared to late treatment (***Muthuri et al., 2014***), while the hazard rate increased with every one-day delay (***Muthuri et al., 2014; Nguyen-Van-Tam et al., 2014***). Prophylaxis with NAIs was shown to be 68%-90% effective in preventing infection (***Kamali and Holodniy, 2013; Ison, 2013***), with low doses leading to lower efficacies and increased emergence of resistance (***Canini et al., 2014***). As a prophylactic measure, using NAIs daily for 6 weeks during an influenza activity period prevented new infections (***McClellan and Perry, 2001***). Early administration—within 48 hours—of NAIs may reduce the risk of illness in close contacts of infected persons (***Nguyen-Van-Tam et al., 2014***).

From the available evidence, it can be seen that NAIs require a strict and narrow time window for small treatment effects to be achieved and, in order to have prophylactic effects, healthy individuals need to take the medicine daily for a long period. This is undoubtedly debatable. The review of ***Jefferson et al***. (***2014***) has suggested that no clinical trials provide concrete evidence for patients, clinicians or policy-makers to use NAIs in annual and pandemic influenza. Furthermore, prophylactic use was also questionable because virus culture was not performed on all trial participants. Therefore, it is not clear whether this is because participants were not infected or because they had an asymptomatic infection ***Jefferson and Doshi*** (***2014***).

Here, we attempt to clarify these claims both mathematically and computationally. Using a within-host infection model of influenza infection, we evaluate the effectiveness of NAIs in reducing viral load and symptom severity as a function of the initiation time of post-infection treatment. Furthermore, using a contact network model of epidemics, we assess the prophylactic effects of NAIs in a population, and discuss treatment strategies with a focus on the cost and availability of the drugs. Our numerical analysis employs oseltamivir as a case study; however, our results and their implications are applicable to NAIs in general.

## Results

### NAIs are unlikely to attain high efficacies even in a best-case scenario

Assuming an idealistic case scenario of instantaneous absorption of NAIs by a treated host as described in Materials and Methods-Eq. (12), the *quasi-steady states* of the drug concentration are given by (details in Appendix 1)

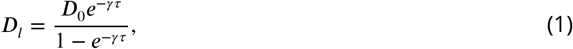

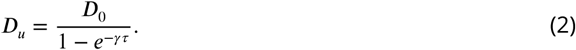

In other words, the drug concentration stabilises to well-defined values after a few doses: an upper bound *D_u_* and a lower bound *D_l_* respectively (Appendix 1). For a given drug and a given treatment regimen, the value of *D_u_* represents a best-case scenario for the therapy. That is, the simplified system defined by Eqs. (7)-(11), supplemented with

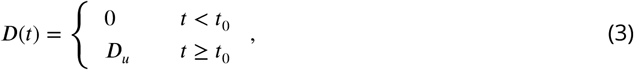

will consistently outperform the full system in terms of the effectiveness of the therapy.

The time-dependent drug efficacy will be given by the reduction in the effective viral replication rate. Denoting this efficacy by *ε* we have, from Eq. (10),

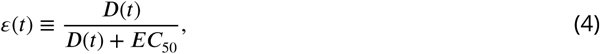

where we immediately see that *D = EC*_50_ results in *ε =* 0.5. From the expression above, it follows that the peak drug concentration *D_u_* translates into a peak efficacy, which we call *ε*^+^, that is given by *ε*^+^ *= D_u_/*(*D_u_ + EC*_50_). Thus, given a particular drug with an elimination rate *γ* and *EC*_50_, we can find the correct values for the dose and administration interval—*i.e*., the treatment regimen—that yield a desired peak efficacy. Introducing Eq. (2) into the expression for *ε*^+^, we obtain the following relationship between *D*_0_ and *τ*:

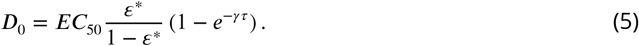

This relationship is illustrated in Fig. 1, which shows contour lines of constant *ε*^+^ as a function of τ and *D*_0_ for the parameter values corresponding to oseltamivir (Table 1). We can observe that the efficacy landscape varies wildly depending on the exact value of *EC*_50_. However, even for an optimistic choice of this parameter—in the case shown, *EC*_50_ *=* 5 mg (middle panel)—reaching a high *ε*^+^ requires very large doses or very frequent intakes. From Eq. (5) we find that, for a peak efficacy of *ε*^+^ *=* 0.99 to be attained, a fixed dose of *D*_0_ *=* 150 mg—corresponding to the pandemic dose for oseltamivir—need be administered approximately 10 times per day, while if we fix the frequency to, *e.g*., 4 times per day, a dose of ca. 275 mg would be required. In this case, *EC*_50_ *=* 5 mg, we note that the values of *ε*^+^ for oseltamivir in the curative and pandemic regimens are given, respectively, by *ε*^+^ ≈ 0.95 and *ε*^+^ ≈ 0.97.

**Figure 1.**
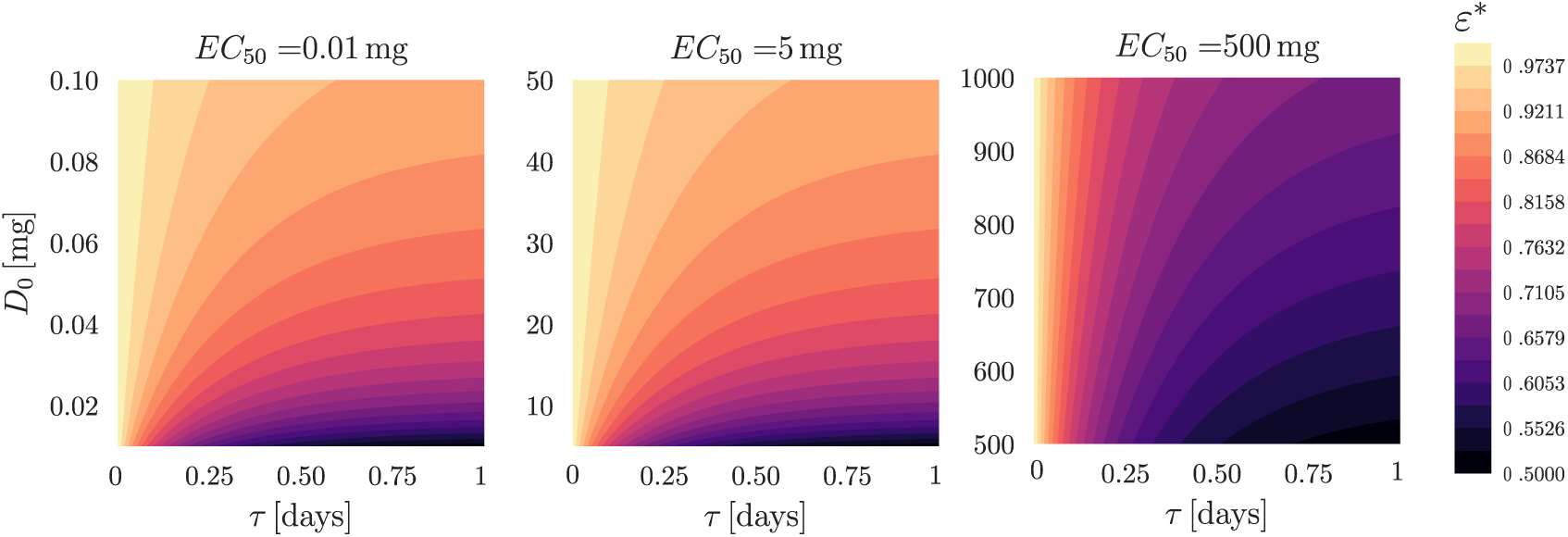
Contour lines of constant peak efficacy *ε*^+^ as a function of the treatment regimen, Eq. (5), with drug parameters from Table 1.

**Table 1.** Parameter values of the within-host model, Eqs. (7)-(12) (***Lukens et.al., 2014***). The initial conditions correspond to a completely susceptible target cell population, *i.e., T*_0_ = 1 and an inoculum size *V*_0_ = 7.5 × 10^−6^TCID_50_ mL^-1^.

| Parameter | Value | Unit | Note |
| --- | --- | --- | --- |
| $\beta$ | 0.0674 | $\text{TCID}_{50}^{-1} \text{ mL day}^{-1}$ | |
| $k$ | 3.684 | $\text{day}^{-1}$ | |
| $\delta$ | 1.364 | $\text{day}^{-1}$ | |
| $p$ | 40356 | $\text{TCID}_{50} \text{ mL}^{-1} \text{ day}^{-1}$ | |
| $c$ | 8.0 | $\text{day}^{-1}$ | |
| $\theta$ | 2.75 | $\text{day}^{-1} \text{ S}$ | S: the symptoms score units |
| $a$ | 0.498 | $\text{day}^{-1}$ | |
| $\gamma$ | 3.26 | $\text{day}^{-1}$ | |
| $EC_{50}$ | 0.01–500 | mg | see Materials and Methods |
| $D_0$ | 75 (c), 150 (p) | mg | The curative (c) & pandemic (p) treatment |
| $\tau$ | 0.5 | day | |

A clearer picture may be obtained by expressing the equation above as a relationship between the drug parameters themselves, fixing the form of the therapy instead. This removes the dependence of the landscape on the half-maximal concentration of a particular drug, and helps shed light on the behaviour of different NAIs. This relationship reads

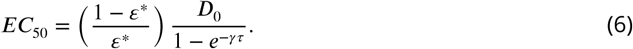

Figure 2 shows the landscape of peak efficacies for a given drug, assuming that the therapy follows either the curative or the pandemic regimen of oseltamivir. Here, for the pandemic regimen, we find that in order to achieve a peak efficacy of at least *ε*^+^ = 0.95, a drug with, *e.g*., a unit elimination rate (*γ* = 1 days^-1^) would need to have a half-maximal concentration no larger than ca. 20 mg. In the curative regimen, in turn, this upper limit goes down to about 10 mg. If we take again the range of reported *EC*_50_-values for oseltamivir as an example in this hypothetical case of a drug that is slowly cleared, we see that the great majority of these concentrations fall below a peak efficacy of *ε*^+^ = 0.95—over 90% if we assume they are uniformly distributed.

**Figure 2.**
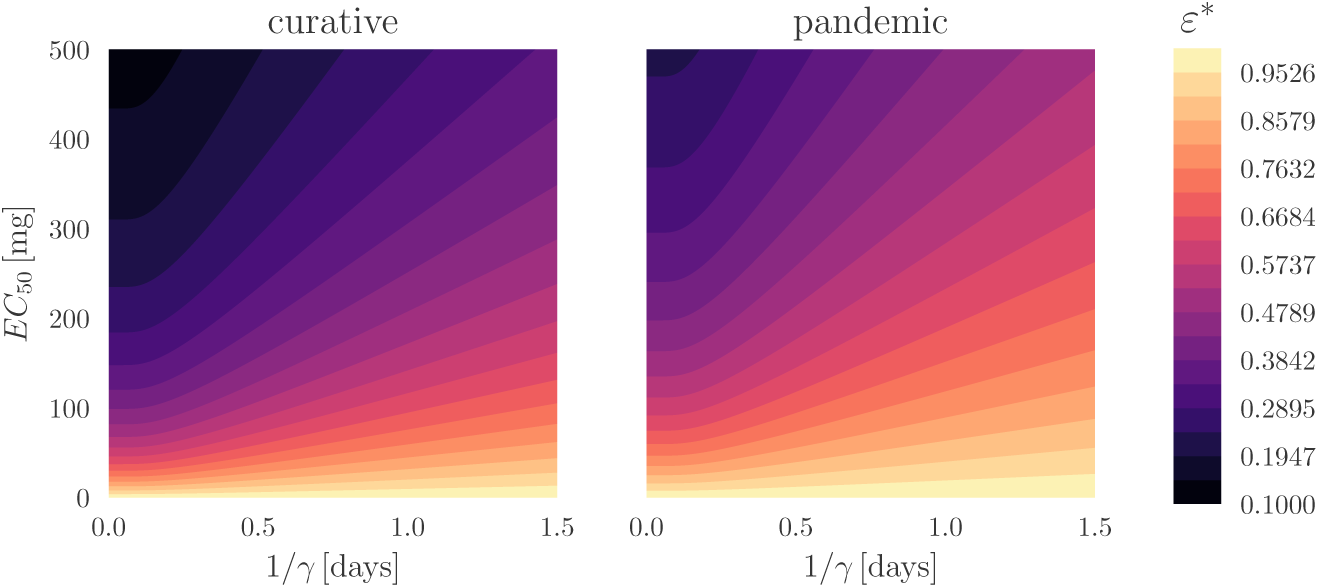
Contour lines of constant peak efficacy ε^+^ as a function of the NAI, Eq. (6). Left: curative regimen for oseltamivir, *D*_0_ *=* 75 mg, *τ* = 0.5 days; right: pandemic regimen for oseltamivir, *D*_0_ *=* 150 mg, *τ* = 0.5 days.

### Even with high efficacy, NAIs effects are negligible in practical settings

The top panels of Figure 3 show the fraction of reduction in the viral load and symptoms AUC in the best-case scenario (*ε*(*t*) = *ε*^+^), which we denote by *χ*, with

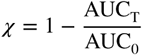

where AUC_T_ corresponds to the case with treatment and AUC_0_ to the base case without treatment. Results are shown for different starting times of the therapy, *t*_0_, in the curative regimen of oseltamivir. The bottom panels of Fig. 3, in turn, show the temporal dynamics of *V* and Ψ under the same conditions.

**Figure 3.**
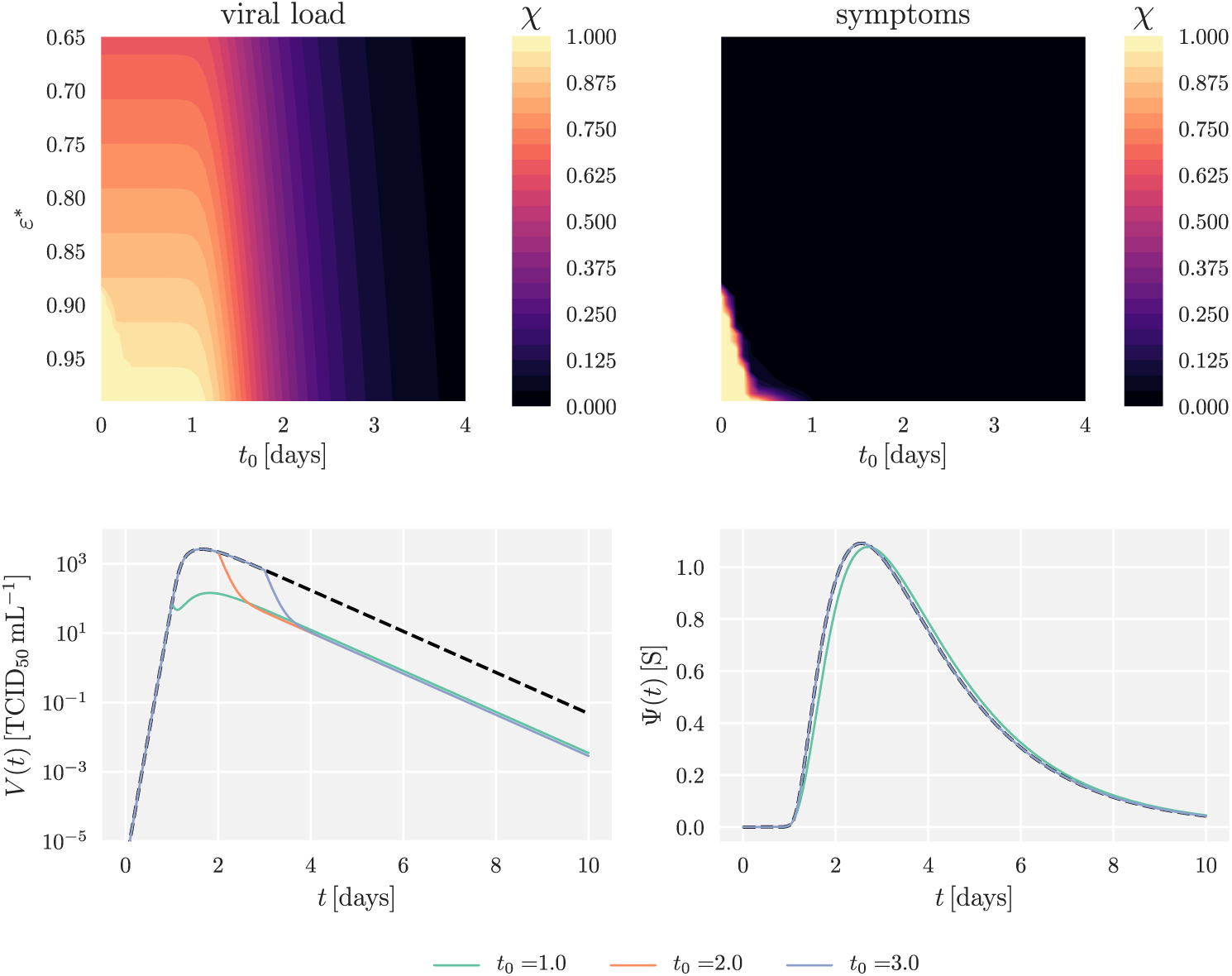
**Top:** Heatmaps showing the fraction of reduction in AUC, *χ*, as a function of treatment initiation times and peak drug efficacy; **left:** reduction in viral load; **right:** reduction in symptoms scores. **Bottom:** Dynamics of the within-host system in the case without treatment (black, dashed lines) and in the case with treatment for ε^+^ = 0.95 and different therapy starting times (coloured lines); **left:** viral load *V*; **right:** symptoms score Ψ. In order to avoid artificial growth of the virus late after infection, simulations with a viral load below a prescribed tolerance—in this case, 10^−3^ TCID_50_ mL^-1^– one day post infection are assumed stop growing, *i.e*., the right hand side of Eq. (10) becomes -*cV* from day 1 onwards for these cases. In all cases, the treatment corresponds to the curative regimen for oseltamivir: *D*_0_ = 75 mg, *τ* = 0.5 days.

Simulations show that even for very large values of *ε*^+^, there exists an optimal starting time for the therapy, and we can appreciate that a late start of the treatment has little to no effect on the dynamics of the infection. In a real-life scenario, it is highly unlikely that the therapy will start during the optimal time window: treatment would not start without symptoms, and seasonal influenza has an incubation period of approximately 2 days (***WHO, 2016***).

Furthermore, we note that, even if there is a substantial reduction in viral load for early starting times, this does not translate into an appreciable effect on the severity of the symptoms. In the case portrayed, a treatment starting as early as 48 h post infection (*t*_0_ = 2 days) results in behaviour which is virtually indistinguishable from the case of an infection without treatment.

### Modest prophylaxis effect on epidemic size: coverage and duration trade-off

Epidemic simulations with one initially infected case provided an average reproductive number of 2.1, resembling previous estimates for seasonal and pandemic flu (***Coburn et al., 2009; Mills et al., 2004***). Upon reproducing this property, different combinations of coverage and duration of the antiviral therapy were implemented, as shown on the left panel of Fig. 4. The durations were chosen based on literature recommendations for responding to influenza outbreaks (***Smith, 2010; Ward et al., 2005***), whereas the coverage fractions were chosen arbitrarily.

**Figure 4.**
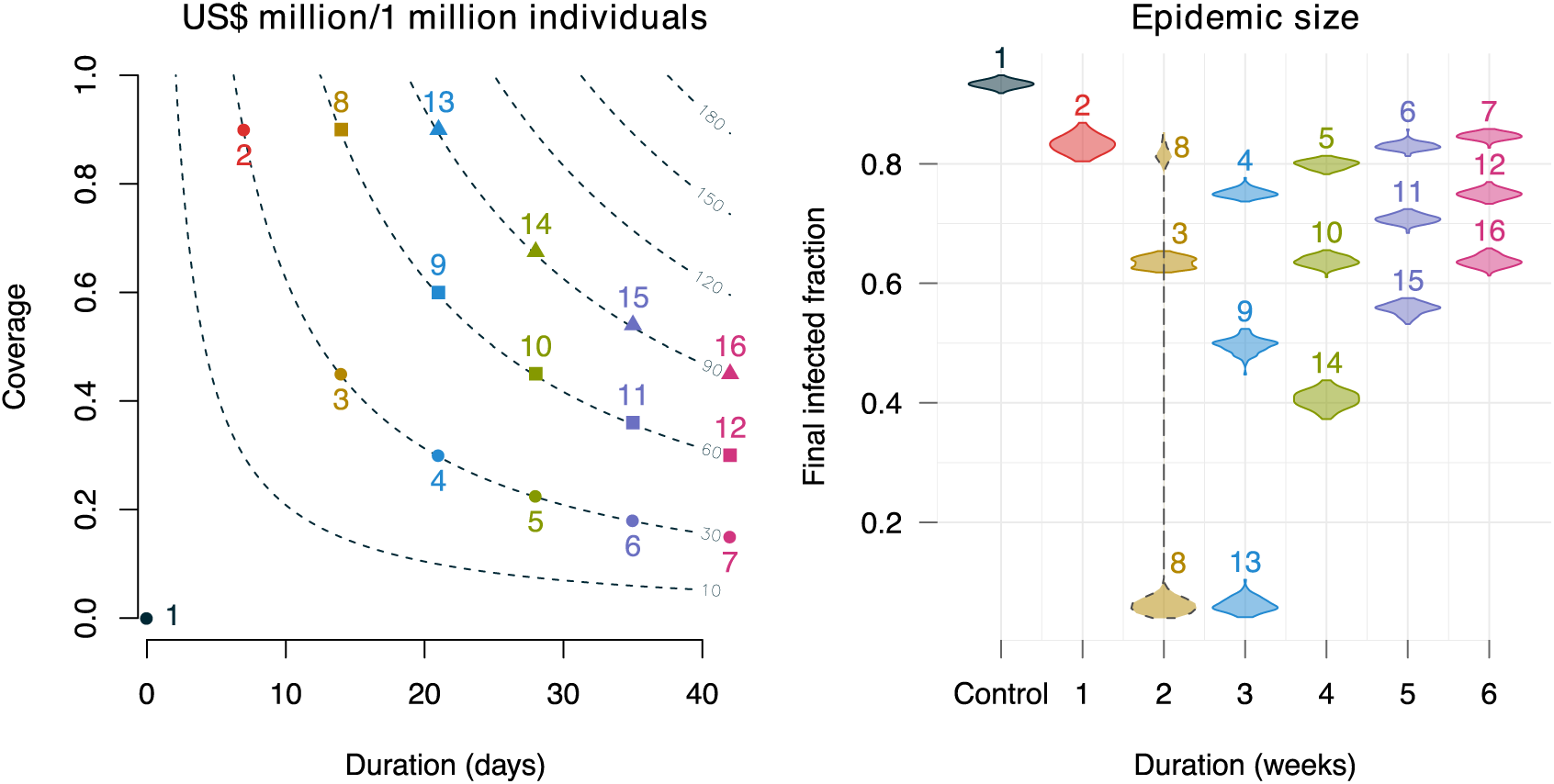
Effects of using oseltamivir on the epidemic size, for different duration and coverage during an epidemic period. **Left:** Total cost per person for the selected durations and coverages. The contour lines show the cost in U.S. dollar per individual, assuming the pandemic regimen (150mg, twice daily) with 0.16 U.S. cents per mg (***Enserink, 2006***). The numbers indicate the simulated scenarios. Three levels of investments are assumed, from 30 to 90 million dollars per a population size of 1 million. **Right:** Violin plots of the final epidemic size for the corresponding scenarios in the left panel. Each violin corresponds to an average of 100 runs. The densities are calculated with non-parametric density estimation using the Sheather-Jones method (***Sheather and Jones, 1991***). Note that the epidemic size distribution in scenario 8 is bimodal.

The right panel of Fig. 4 shows that that there were reductions in epidemic size, but in many cases the reduction was small given the investment. In scenario 7, for example, 30 million dollars were spent over 6 weeks but the epidemic size reduction was lower than 20% compared to the case without intervention (scenario 1). Tripling the allocated resources (scenario 16) brought the epidemic size further down only by ca. 25%.

In addition, the right panel of Fig. 4 also shows that there was a trade-off between coverage and duration. Generally, prolonging the duration while keeping a low coverage was not efficient (scenarios 5–7, 10–12, and 14–16). However, a very high coverage with a short duration (scenario 2) was not useful either. There seemed to be a bifurcation point between two and three weeks where the scenarios tended to converge. Scenarios 8 and 13 illustrate this phenomenon: both were ideally covered (90%), but in one of them (scenario 8) some of the simulations resulted in large epidemics, while the other provided a complete control of the epidemic.

## Discussion

Neuraminidase inhibitors (NAIs) constitute the primary type of antiviral drugs against influenza. Their effectiveness in reducing the spread of the virus depends strongly on the dosage and interval between intakes, and is dramatically hindered by a late start of the treatment. We have evaluated the within- and between-host effects of treatment with NAIs as a function of the time of initiation, using a constant-concentration model for the drug. ***Palmer et al***. (***2017***) introduced a constant-concentration model that ***approximates*** the resulting viral load from the full model time-dependent under certain conditions. Since our aim is to define lower and upper bounds for efficacy, our approach has been slightly different: we have derived the ***exact*** values to which the drug concentration quickly converges at its least and most effective, and have used the latter as a best-case scenario for the therapy.

Our analytical results suggest that, even if we assume that the drug reaches its peak concentration at the time of initiation of the treatment, and remains at this constant value from that moment onwards, it is unlikely that an extremely large reduction in viral replication rate will be achieved under realistic conditions. In order for this to occur for typical dosages and intake frequencies, the drug must have a very low *EC*_50_ value. As an example of this, most of the range of half-maximal concentrations reported for oseltamivir result in peak efficacies which are below 95%, even if we consider a mean life of one day for the drug. We further note that the volumes from ***Rayner et al***. (***2008***) were obtained for oseltamivir in combination with probenecid, a potent competitive inhibitor of the renal tubular secretion of weak organic acids ***Howton*** (***2006***). This co-administration has a pronounced effect on oseltamivir carboxylate pharmacokinetics, reducing renal clearance and increasing the area under the concentration-time curve (AUC) (***Howton, 2006; Davies, 2010; Wattanagoon et al., 2009***). The apparent volume of distribution for oseltamivir alone administered orally can greatly surpass 100 L (***Grayson et al., 2017***), which translates into the values we have used as upper and lower bounds for the half-maximal concentrations—shown in Table 1—being as least twice as large, with the corresponding decrease in peak efficacy, as can be observed from Fig. 2.

Numerical simulations of the within-host system showed that, even for very large values of the peak efficacy, the reduction in viral replication rate does not induce a similar impairment on the severity of the symptoms. While this is a direct consequence of the latter being directly dependent on the proportion of infected cells, rather than on the viral load—see the form of Eq. (11)—we believe that this choice is well justified in ***Lukens et al***. (***2014***). Furthermore, even in the case of the viral load itself, an appreciable reduction is only appreciable for a very early start of the treatment. In reality, the starting time for the therapy will usually lie somewhere around the start of the third day post infection, and the best outcomes observed from the model are unlikely to occur in a real-world setting.

Embedding the within-host dynamics into an epidemic model of influenza transmission, we assessed the level of protection conferred by NAIs and their effectiveness against an outbreak when taken as a prophylactic measure. Given a limited availability of the drug as well as resources in the affected countries, selection of prevention strategies need be well-informed for cost-effectiveness. We found that there was a trade-off between duration and coverage and it seemed that prolonging coverage over three weeks is not cost-effective. A best-case scenario corresponds to providing the drug for more than two weeks with as high a coverage as possible. However, here we observed some cost-effective strategies only in highly favourable conditions; in reality, there are many conditions that would further compromise the effect of the drug: (i) There can be repeated introduction of newly infected cases into the population, e.g., from travelling, immigration; this can make the short coverage duration ineffective; (ii) The actual drug stockpiles in the 2001 H1N1 pandemic ranged only from 0.1% to 25% in rare cases (***Meave Gutierrez-Mendoza et al., 2012***). With many affected countries having large populations, there could be severe shortages in supply. Nevertheless, in a small community, these estimates of required coverage and duration could hold given that routine public health practices for influenza epidemic are in-place, e.g., examining newcomers, providing quarantine and isolation measures, closing schools and social activities. But the effect of the drug in this case can be very difficult to asses.

It is important to remark that, by turning to a description based on the peak efficacy for a given treatment regimen, we have rendered the analysis essentially independent of the drug parameters in the sense that only the actual value of the peak efficacy from Eq. (6) will depend on them, but their functional relationship and the landscape represented in Fig. 2 will not. Therefore, while we have carried out our analysis using oseltamivir as a case study, we expect our results to hold for NAIs in general. We also stress that all of the analysis above has considered the ***best possible case***, and that the impact of the therapy in a practical scenario will be lower than observed here due to the fluctuating nature of the efficacy and the aforementioned late starting times in the case of post-infection therapy.

Taken together, our results imply that the use of NAIs is only warranted as a prophylactic measure on very limited conditions, and formalise the claims that their use in therapeutic situations will result in impaired performance in most real-case scenarios. Interestingly, it may even be detrimental as influenza virus can mutate and reassort to circumvent available drugs such as oseltamivir and zanamivir (***Hurt et al., 2009***). Therefore, NAIs should be prudently used to avoid the development of drug-resistant strains (H275Y and I223R) ensuring they remain an effective defence against future lethal influenza viruses.

## Materials and Methods

### Within-host infection dynamics

The within-host model of the dynamics of influenza infection corresponds to the target cell-limited model with delayed virus production originally introduced by ***Baccam et al***. (***2006***), and later extended by ***Lukens et al***. (***2014***) in order to consider the effects of the virus on individuals' symptoms. Here, we extend the model further to take into account the effects of treatment with NAIs. The full system of equations is given by

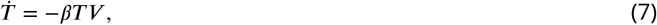

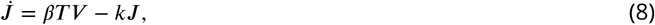

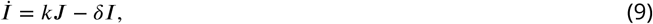

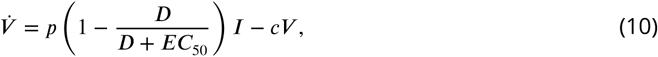

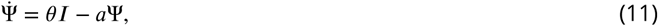

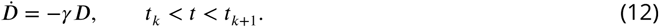

The system considers a population of target (epithelial) cells, divided into susceptible (*T*), infected (*J*) and productively infected (*I*). After infection with the virus, occurring at a rate *β*, susceptible cells enter the latent phase *J*, where they remain for an exponentially-distributed time with mean 1/*k*, after which they enter the productively infected class *I*. Infected cells shed virus at a rate *p* and have a mean lifespan 1/*δ*. The free virus, in turn, is cleared at a constant rate *c*. The intensity of the symptoms, denoted by Ψ, increases with the proportion of infected cells at rate *θ* and has a constant decay rate a.

We include the effect of the treatment with NAIs as a reduction factor in the rate of virus shedding *p*, which increases with the drug concentration *D* in a sigmoid fashion. The drug itself is assumed to be eliminated at a rate *γ*. The different *t_k_*, with *k* = 0, 1, 2,…, represent the times of drug intake. We take constant administration intervals *t_k_*_+1_ *− t_k_ ≡ τ*, and a constant dose equal to *D*_0_. These two parameters, *τ* and *D*_0_, define the treatment regimen; the elimination rate *γ* and half-maximal concentration *EC*_50_ —that is, the concentration at which the drug reaches a 50% efficacy—constitute the relevant drug-dependent parameters. We note that this is a simplified model for the NAIs, and a more accurate description of the dynamics of the system would require consideration of the drug's absorption and conversion into its active metabolite (*Canini et al., 2014*). Here, we essentially equate the concentration of the latter with D. Since we are interested in finding an upper bound for constant-concentration efficacy, this simplified model is sufficient for our analysis.

The parameter values for Eqs. (7)—(11) are taken directly from ***Lukens et al***. (**2014**). We start by focusing on the effects of therapy with oseltamivir, and therefore we consider an elimination rate *γ* = 3.26 days^-1^ (***Wattanagoon et al., 2009***) and a half-maximal concentration ranging from ca. *EC*_50_ = 0.0008 *μ*M to ca. *EC*_50_ = 35 *μ*M, with 1 *μ*M ≈ 0.284 mg/mL (***Tamiflu, 2009***). Considering an apparent volume of distribution for oral administration of ca. 50 L (***Rayner et al., 2008***), this translates into a range from ca. *EC*_50_ = 0.01 mg to ca. *EC*_50_ = 500 mg, which is what we use in our analysis. The parameters of the treatment itself correspond to the curative and pandemic regimens of oseltamivir: respectively, *D*_0_ = 75 mg and *D*_0_ = 150 mg, in both cases administered twice a day (*τ* = 0.5 days) (***Canini et al., 2014***). All parameters are listed in Table 1. The Python code for this section can be visited at @ systemsmedicine/neuraminidase-inhibitors.

### Epidemic simulations

To assess the prophylactic effects of NAIs in an epidemic context, the model defined by Eqs. (7)-(12) was used to generate the infection dynamics of an individual-based network model of influenza transmission. Epidemic settings were tailored to detect the drug's effect by homogenising the scenarios as follows: (a) All infected individuals have the same disease progression and respond similarly to the drug; (b) Uninfected individuals are equally susceptible to the infection without immunity, i.e., transmission is defined only by the infecter; (c) The drugs are readily available and can be delivered to all intended recipients uniformly; (d) All intended recipients take the drugs with complete adherence; (e) All infected cases are reported, including asymptomatic cases; (f) There are no other interventions in-place and the contact network remains unchanged during the epidemic. In this way, changes in epidemic trajectories can be attributed solely to the drug's effect.

We considered transmission conditions similar to those in ***Lukens et al***. (***2014***), i.e.: (i) The transmission potential of an infected subject *i* at any given time is defined by its viral load at that time, normalized by the maximum viral load, i.e., *p_i_*(*t*) = *V_i_*(*t*)/max(*V*)—noting that this depends on both the time of contact and the time since infected; (ii) The infectious period is started when the viral load crosses the threshold *V_c_* = 1.35 to ensure the incubation period and infectious period conform to clinical observations.

For the population model, we used a simulated static network with a size of 10,000 nodes that embodies the average contact distribution and contact patterns of ten European countries (***Mossong et al., 2008***). Simulations were then carried out as follows: (1) Seed 100 random infected nodes (note that to calculate the basic reproduction number, only one infected node is seeded); (2) Check connected nodes of the infected cases and evaluate Bernoulli trials with probabilities of success *p_i_*(*t*)*, i = 0, …,I*(t), where *i* and *I*(*t*) denote infected node *i* and the total number of infected nodes at time t, respectively; (3) Move to the next time step and repeat step (2) until *I*(*t*) *=* 0. For computational efficiency, we ran the simulations with a one-day time step and added random noises to the time of infection by sampling from a uniform distribution *U*(−.5, .5), representing different times of contact during a given day.

Simulated scenarios were assumed constrained by a fixed amount of resources (U.S. dollars) calculated based on the epidemic regimen of 150mg twice daily, and the minimum price for oseltamivir in large purchases: 1.6 U.S. cents per mg as of 2006 (***Enserink, 2006***). Based on a given investment, scenarios can be differed by the proportion of the population to be covered and the time during which uninfected subjects within coverage can be provided with the intended amount of drug without any disruptions. Each scenario was simulated 100 times to obtain distributional epidemic trajectories.

Epidemic simulations were written in R language (***R Core Team, 2015***) and run on the FUCHS cluster operated by the Center for Scientific Computing (CSC) of the Goethe University Frankfurt. The R code for this section can be visited at @ systemsmedicine/neuraminidase-inhibitors.

## Acknowledgements

The authors would like to thank the Center for Scientific Computing (CSC) of the Goethe University Frankfurt for facilitating the FUCHS cluster for the simulations. This work was supported by the Alfons und Gertrud Kassel-Stiftung.

## Author Contributions

C. P.-R. constructed the within-host model and performed the simulations. V. K. N. constructed the multiscale model and performed the simulations. G. H.-M contributed with the PK/PD model and simulations. E. H.-V. envisaged and supervised the project. All authors discussed and wrote the paper.

## Appendix 1

### Quasi-steady state of the drug concentration

**Appendix 1 Figure 1.**
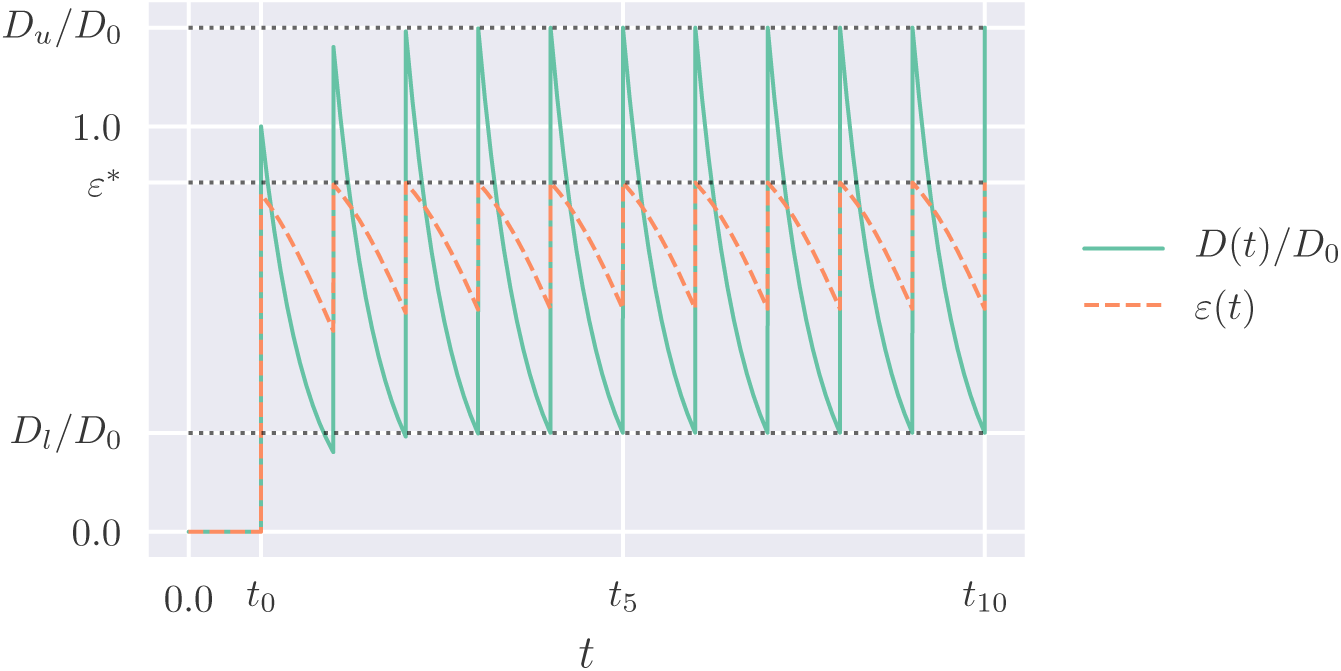
Green line: illustration of the drug dynamics when administrated at constant time intervals and fixed dose *D*_0_, starting at *t = t*_0_; orange, dashed line: the corresponding drug efficacy, Eq. (4). The grey, dotted lines signal the lower and upper bounds for the drug concentration after the quasi-steady state is reached, respectively Eqs. (1) and (2), as well as the peak efficacy *ε*^+^.

We can find the quasi-steady states illustrated in Appendix 1 -Fig. 1 by noting that, when we start the therapy,

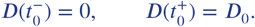

After a time *τ*, the initial value of *D* will have decayed by an amount equal to *e*^-^*^γτ^*, so that the drug concentration just before and just after the second administration will be given by

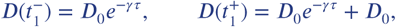

where we have used that the interval between intakes is constant and equal to *τ*. Repeating this reasoning a few more times we find that, when we reach time *τ_n_*, the drug concentration has the form

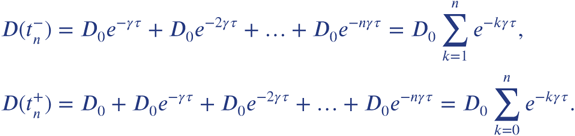

The summations in the expressions above have a closed form when *n* → ∞, provided that +*γτ > 0* (***Abramowitz and Stegun, 1965***), which is naturally the case since both parameters are positive. Therefore, we can obtain the quasi-steady state values as

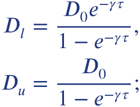

these are reproduced in Eqs. (1)-(2) of the main text.

A similar approach can be used to obtain the values to which the peaks and valleys of the drug concentration converge after a long time has passed for the more complicated model found in ***Palmer et al***. (***2017***). These lower and upper bounds are given by

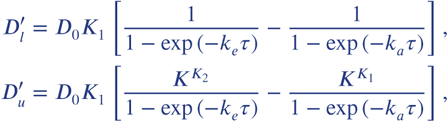

where *k_a_* and *k_e_* are, respectively, the absorption and elimination rates of the drug, and

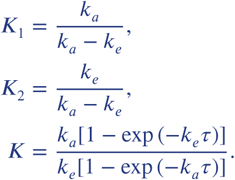

We find that 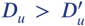, and therefore *D_u_* may be safely used as an upper bound for a best-case scenario.

